# Strengthening Phage Resistance of *Streptococcus thermophilus* by Leveraging Complementary Defense Systems

**DOI:** 10.1101/2024.10.22.618286

**Authors:** Audrey Leprince, Justine Lefrançois, Damian Magill, Philippe Horvath, Dennis A Romero, Geneviève M. Rousseau, Sylvain Moineau

## Abstract

CRISPR-Cas and restriction-modification systems represent the core defense arsenal in *Streptococcus thermophilus* to block lytic phages, but their effectiveness is compromised by phages encoding anti-CRISPR proteins (ACRs) and other counter-defense strategies. Here, we explored the resistome of 263 *S. thermophilus* strains to uncover other anti-phage systems. The defense landscape of *S. thermophilus* was enriched by 21 accessory defense systems, 13 of which had not been previously investigated in this species. Experimental validation of 17 systems with 14 phages showed varying anti-phage levels, uncovering intra-genus specificities among the five viral genera infecting *S. thermophilus*. Interestingly, the resistance levels were even higher when some defense systems (Dodola and PD-Lambda-1) were expressed from a low-copy plasmid or when integrated into the chromosome. We also observed a synergistic effect when combining Gabija with CRISPR-Cas, underscoring the potential of these additional defense systems for developing more robust industrial *S. thermophilus* strains, particularly against ACR-encoding phages.

## 1. Introduction

*Streptococcus thermophilus* is a lactic acid bacterium (LAB) widely used to produce fermented dairy products such as yogurt and certain cheeses. In the dairy industry, large-scale milk fermentation processes are meticulously controlled to ensure the quality and consistency of the final product. However, phage infections present a significant risk for these added bacterial cultures as they can reduce bacterial growth and milk acidification rates, thereby disrupting the fermentation process^1^. These delays or halts in fermentation can result in poor-quality products, waste, and economic losses. Therefore, controlling virulent phages in this environment is paramount and requires constant monitoring, explaining why phages that infect LAB, including *S. thermophilus,* have received significant research attention^2^. Because phages are impossible to eliminate from manufacturing facilities due to constant entry through milk supply, the dairy industry relies on a variety of control strategies to mitigate phage contamination and dissemination^3^. One crucial approach is using industrial strains with increased resistance against known circulating phages^2^. To date, five viral genera are known to infect *S. thermophilus*, with phages belonging to the *Moineauvirus* genus (formerly *cos* group) and the *Brussowvirus* genus (formerly *pac* group) being the most predominant, isolated in 69% and 29% of cases, respectively^2^. Members of the three other genera, *Vansinderenvirus* (formerly 5093 group), *Piorkowskivirus* (formerly 987 group), and P738 are more rarely isolated.

Bacteria have evolved multiple defense systems to counter phage infection at different stages of the viral replication cycle^4^. In *S. thermophilus* strains, Clustered Regularly Interspaced Short Palindromic Repeats and associated protein (CRISPR-Cas) systems are highly prevalent and have been well characterized^5,6^. By challenging sensitive strains with problematic phages, strains that acquire additional spacers and exhibit increased phage resistance can be selected and then used in starter culture blends or included in strain rotation strategies^7^. In addition, this relatively minor genomic modification does not affect the strain’s other technological properties. As this adaptive immunity system can naturally improve the phage resistance profile, the derived industrial cultures have not required regulatory approval. Thus, CRISPR-Cas systems have become fundamental in generating bacteriophage-insensitive mutants (BIMs) that are now used worldwide^7,8^.

However, due to the constant evolutionary race with their host, phages have developed counter-defense mechanisms to bypass CRISPR-Cas immunity^9^. For instance, CRISPR escape mutants (CEMs) have acquired mutations in their protospacer or protospacer adjacent motif (PAM) to prevent matching with their host spacer, thereby abolishing interference^10,11^. In addition, several *S. thermophilus* phages encode one or multiple anti-CRISPR (ACR) proteins that neutralize Cas9, thereby preventing DNA binding or nuclease activity^12–14^. Therefore, it is crucial to explore alternative strategies for generating phage-resistant strains to safeguard CRISPR-Cas defense.

The diversity of bacterial anti-phage defense systems has been extensively explored in recent years, uncovering over 150 distinct systems with diverse mechanisms from various bacterial species^15,16^. In *S. thermophilus*, although research has primarily focused on CRISPR-Cas systems, early studies also characterized several restriction-modification (RM) systems^17–19^ as well as a few prophage-encoded lipoproteins, the latter protecting against related phages through superinfection exclusion^20,21^. More recently, Kelleher *et al*.^22^, while investigating the methylome of 27 industrial *S. thermophilus* strains, highlighted the significant role RM systems play in defense in this species. They also identified nine additional defense systems in these strains, with seven of them (AbiD, AbiE, Gao19, Gabija, Hachiman, Kiwa, and SoFic) conferring low-to-moderate resistance against four streptococcal phages while two did not (Dodola, PD-T4-6)^22^. Through the analysis of other *S. thermophilus* genomes, they also identified, but not tested, five other defense systems (AbiH, Borvo, PD-Lambda-1, PrrC, and RloC)^22^.

In this study, we significantly advanced our understanding of the *S. thermophilus* resistome by systematically analyzing known defense systems in 263 publicly available genomes and investigating counter defense strategies in streptococcal phages. We identified 28 defense systems, confirming that CRISPR-Cas (three types) and RM systems (four types) function as core lines of defense. However, our findings also revealed that phages have developed multiple counter-defense strategies to bypass these systems, highlighting the need for a deeper investigation of other defense mechanisms. Among the 21 other defense systems identified, we showed effectiveness of 17 of them against several streptococcal phages and lactococcal phages. Importantly, by combining some of the most effective defense systems with CRISPR-Cas, we were able to enhance the overall resistance levels, especially against phages that encode ACRs.

## 2. Results

### *S. thermophilus* resistome expands well beyond CRISPR-Cas and RM systems

Defense systems were predicted in publicly available genomes of *S. thermophilus* (N=263, as of March 2023) using DefenseFinder^23^ and PADLOC^24^ (Fig. 1A and Supplementary Table S1-S2). While both tools generally provided similar predictions, we noticed that each missed some defense systems (Supplementary Table S2). For example, type IV RM systems were only detected by PADLOC, while AbiH was only identified by DefenseFinder, despite both bioinformatic tools containing models for these systems. By combining their outputs, we achieved a more comprehensive analysis across our large dataset, which includes incomplete genomes, enabling the detection of less common defense systems (Fig. 1A). This approach revealed 28 distinct defense systems, significantly expanding the known diversity of the *S. thermophilus* resistome^22^. Among these, we identified three CRISPR-Cas types (II-A, III-A, and I-E), four RM types (I-IV), and 21 additional defense systems, 13 of which were not previously tested in Kelleher *et al*.^22^. The number of defense systems per strain ranged from 3 to 13, with an average of 7.5 systems per strain (Fig. 1B), which is similar to the numbers reported by Kelleher *et al*.^22^. Plasmids and prophages are rare in *S. thermophilus*, detected in only 12.9% (36/263) and 6.8% (18/263) of the strains, respectively (Supplementary Table S1). Consequently, except for four RM systems and one AbiD homolog, all defense systems were chromosomally encoded, and none were found within prophages (Supplementary Table S2). Nonetheless, a comparison of defense system distribution across strain phylogeny revealed that these systems are not only shared among closely related strains but also between strains from different phylogenetic clades, suggesting that horizontal gene transfer mechanisms, notably natural competence which is common in *S. thermophilus*, play a role in their dissemination (Fig. 1C and Supplementary Fig. S1).

**Fig. 1:**
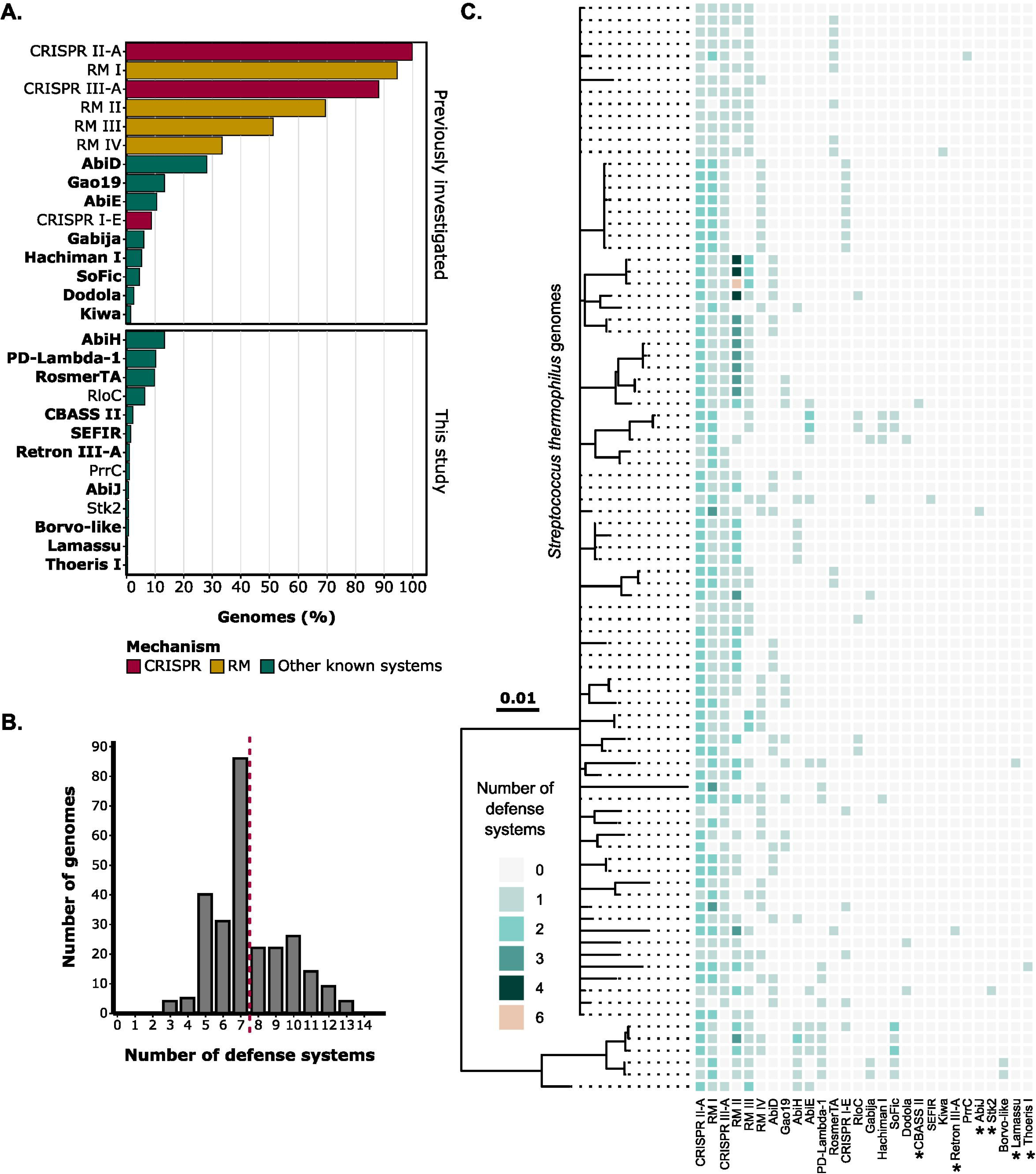
*S. thermophilus* anti-phage defense systems extend beyond CRISPR-Cas and RM systems. **A.** Prevalence of known defense systems in RefSeq genomes of *S. thermophilus* (N=263, as of March 2023). These systems are categorized into previously reported defenses and newly identified ones in this study. The 18 systems experimentally validated in this work are highlighted in bold. The “Borvo-like” system differs from the original Borvo, being annotated as a two-gene system composed of two BovA subunits, rather than one. **B.** Distribution of the number of anti-phage defense systems per genome. The red dashed line indicates the average number of defense systems per *S. thermophilus* genome. **C.** Phylogenetic tree of complete *S. thermophilus* genomes (N=85), illustrating the distribution of identified defense systems. Six incomplete genomes were included to represent defense systems which were not found in complete genomes (marked with an asterisk).

### CRISPR-Cas and RM systems are extensively found in *S. thermophilus* strains

CRISPR-Cas systems have been widely investigated in *S. thermophilus*, due to their prevalence and ability to be easily manipulated for industrial purposes. Previous studies have identified four distinct CRISPR loci (CR1-4)^25,26^. Among these, two well-known type II-A CRISPR-Cas systems associated with the CR1 and CR3 loci, were highly prevalent in *S. thermophilus* strains (Fig. 1A and 2A). Specifically, the CR1 locus was present in all strains, while 70% (184/263) also harbored the CR3 locus (Supplementary Table S3). These two CRISPR loci have been shown to be very effective at acquiring new spacers^27^. The CR2 locus, classified as a subtype III-A system, and was found in 88% (232/263) of the strains. Finally, the CR4 locus, belonging to a type I-E system, was present in only 9% (23/263) of the strains. Although rare, *in vivo* spacer acquisition in the CR2 locus was recently demonstrated for one strain^28^, however no such acquisition was ever observed for CR4^25^. CRISPR loci frequently co-occurred within the same genome, with most strains encoding two (31%, 81/263) or three (59%, 155/263) loci, suggesting a complementary role of these types of CRISPR-Cas systems^29^ (Fig. 2A). Of note, while type II-C have been reported in studies focusing on *in silico* prediction of CRISPR-Cas systems in *S. thermophilus,* our analysis identified only one such instance, which turned out to be a misprediction of a type II-A system, confirming the absence of type II-C in this bacterial species.

**Fig. 2:**
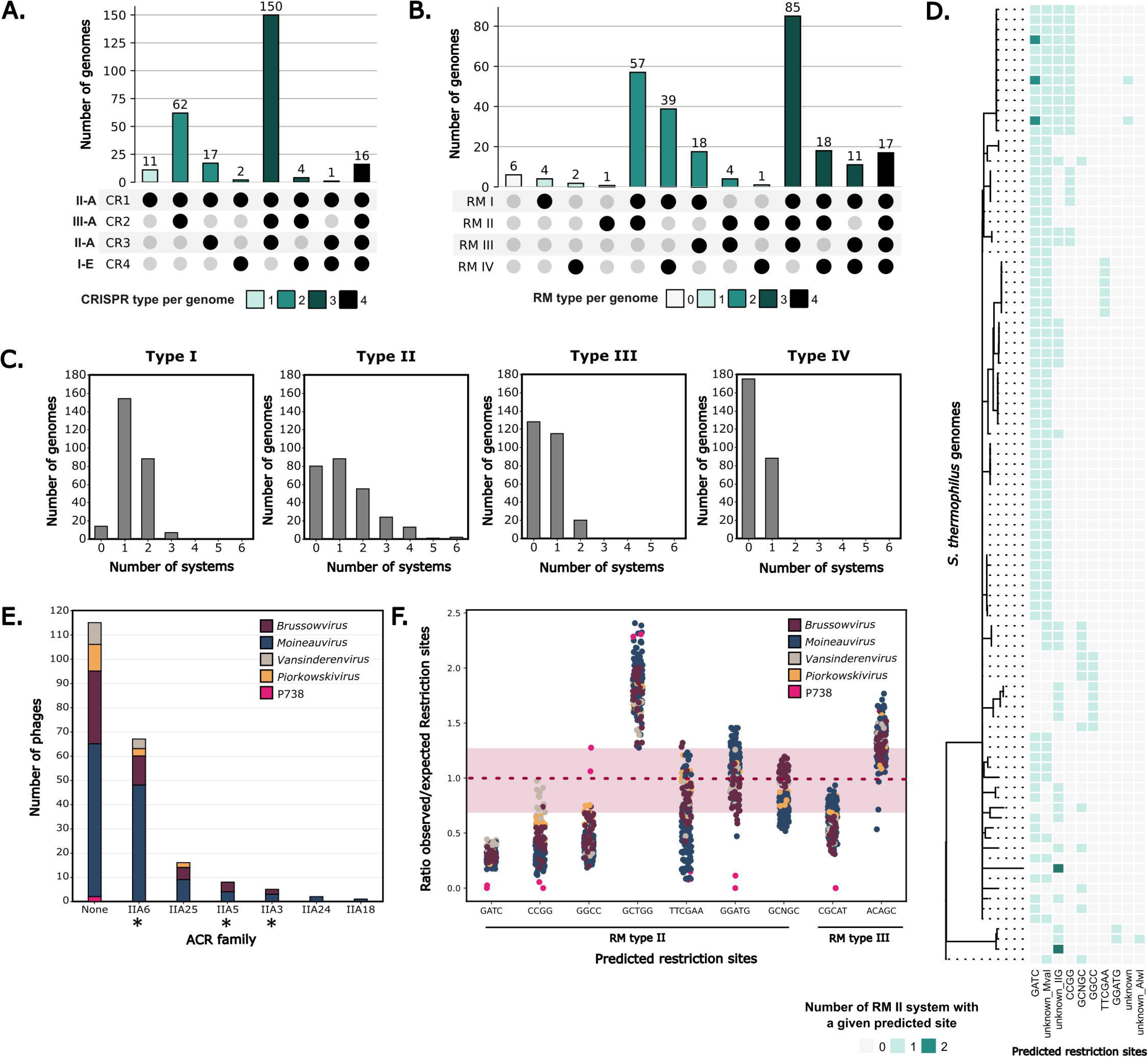
CRISPR and RM systems are widespread in *S. thermophilus,* and phages have several counter defense mechanisms. Co-occurrence of **A.** CRISPR-Cas loci and **B.** RM types within the same *S. thermophilus* genome. **C.** Distribution of the number of RM systems per genome according to the type. **D.** Heatmap of co-occurrence within the same genome of RM type II with different predicted restriction sites. Only strains harboring at least two RM type II systems are shown on the phylogenetic tree (N=95). RM systems with no predicted restriction site were divided into different unknown categories based on the restriction enzyme annotation (MvaII or AlwI) or subtype (IIG). **E.** Prevalence of predicted anti-CRISPR (ACR) proteins in phages infecting *S. thermophilus* (N=191, NCBI June 2023). Experimentally validated ACRs proteins in *S. thermophilus* are marked with a star. Color coding corresponds to phage genus. **F.** Plot of the ratios of observed to expected number of restriction sites in streptococcal phages for RM type II and III restriction sites (predicted with REBASE). The observed number of sites corresponds to the count of each site within the phage genomes, while the expected number was estimated using a Markov-immediate neighbor dependence statistical model^68^. Ratios < 0.75 indicate underrepresentation and ratios > 1.25 indicate overrepresentation.

RM systems were also prevalent in *S. thermophilus*, with at least one RM system present in 98% (257/263) of the analyzed strains, and some harboring up to nine distinct RM systems (Fig. 2B and Supplementary Fig. 2). Type I RM systems were nearly ubiquitous, consistent with the methylome analysis of 27 *S. thermophilus* strains, which attributed most methylated motifs to type I RM, with only a few linked to type II and III RM systems^22^. Type IV RM systems that cleave methylated DNA were found in 33% (88/263) of the strains (Fig. 2C). Like CRISPR-Cas systems, different RM types frequently co-occurred within the same strain (95%, 250/263), likely due to their mode of action, sequence specificities and varying susceptibilities towards phage-encoded anti-RM proteins (Fig. 2B). Multiple instances of the same RM type within a single strain were found for types I, II, and to a lesser extent, III (Fig. 2C). Some strains harbored up to three type I or six type II RM systems. Predictions of putative recognition sites for type II RM systems showed that these co-occurring systems often recognize different sequences, thereby enhancing the strain’s ability to target a broader range of phages with varying restriction site frequencies in their genomes (Fig. 2D and Supplementary Fig. S2B). Similar trends were also observed for type I and III RM systems (Supplementary Fig. S2C-D).

### Streptococcal phages have strategies to counter CRISPR-Cas and RM defenses

Given the widespread presence of CRISPR-Cas and RM systems in *S. thermophilus*, we investigated counter-defense strategies in the publicly available genomes of their infecting phages (N=191). Six families of known ACRs that target type II-A systems were identified in 40% (77/191) of the streptococcal phages (Fig. 2E). Among these, AcrIIA6, which neutralizes the Cas9 nuclease associated with the CR1 locus, was especially frequent. AcrIIA6 was found in 36% (69/191) of the phage genomes, including in 42% (49/116) of the *Moineauvirus* genus, the most prevalent streptococcal viral genus in the dairy industry^2^. AcrIIA5, which inhibits both type II-A CRISPR-Cas systems (CR1 and CR3) and AcrIIA3, which inhibits only the system linked to the CR3 locus, were present in 4.2% (8/191) and 2.6% (5/191) of the phages, respectively. AcrIIA25, AcrIIA24, and AcrIIA18, which have not been characterized yet in *S. thermophilus* phages, were identified in 8.3% (16/191), 1.5% (2/191) and 0.5% (1/191) of the phage genomes analyzed, respectively. Overall, ACRs were more prevalent in phages belonging to the *Moineauvirus* (46%, 53/116) and *Brussowvirus* (35%, 16/46) genera, as compared to phages of the rarely found *Piorkowskivirus* (21%, 3/14), *Vansinderenvirus* (30%, 4/13), and P738 (none).

No known anti-RM proteins was detected in the streptococcal phages analyzed. However, DNA methyltransferases were surprisingly present in 20% of them, indicating that some may evade genome restriction through DNA methylation of recognition sites (Supplementary Table S2). In addition, previous studies have shown that reducing the number of restriction sites within phage genomes increases the likelihood of evading RM^30^. An analysis of restriction site avoidance for the predicted recognition sites of type II and III RM systems in streptococcal phages revealed that certain restriction sites, such as GATC, CCGG, and GGCC, were commonly underrepresented in most phage genomes (Fig. 2F). Others, such as GGATG, were selectively excluded in some viral genera (e.g., P738-like phages). Conversely, the predicted GCTGG site showed no reduction in number across all phages. These data suggest that the reduced numbers of restriction sites seem to be a common evasion strategy against RM systems in streptococcal phages.

### Versatile modes of action in accessory defense systems of *S. thermophilus*

Our bioinformatic analysis uncovered 21 non-CRISPR and non-RM defense systems (Fig. 1A). AbiD was the most prevalent, occurring in 28% (74/263) of the strains. The other systems were present in less than 15% of the strains. Most of them were found as one copy per genome, although a few (AbiD and SoFic) were occasionally present in duplicate within the same genome (Table 1). This co-occurrence suggests a complementary role, potentially targeting different phages and enhancing the global defense of a given strain.

While the mode of action for some of these defense systems remains unknown, others have been characterized in other bacterial species as indicated in Table 1. Four abortive infection (Abi) systems (AbiD, AbiE, AbiH, and AbiJ), originally characterized in the LAB *Lactococcus lactis*, have also been identified in *S. thermophilus*^31^. These systems block phage replication at various stages before cell lysis, thereby preventing the spread of the infection to neighboring cells^32^. Besides the classical Abi systems, most defense systems present in *S. thermophilus* are associated with population-level protection via an Abi phenotype (Table 1). Two key strategies seem to drive Abi defense in *S. thermophilus*: depletion of NAD+ and/or ATP^16,33,34^ within the infected cell (Gao19, Thoeris, and Sefir) and non specific DNA degradation^34–37^ (Gao19, Gabija, Hachiman, and Kiwa). Signal-based defenses like CBASS^38^ and Thoeris^33^, which rely on intracellular signaling molecules (e.g., cyclic oligonucleotides and cyclic ADP-ribose) to activate effector proteins inducing cell death, are also present. Toxin-antitoxin systems^16,39^ (AbiE and RosmerTA), retrons^40^, and phosphorylation of essential cellular pathways by Stk2^41^ are also found in a few strains. Lastly, RloC^42^ and PrrC^43^ systems, which inhibit translation following the neutralization of type I RM systems by phage proteins, were found in 6% (17/263) and 1% (3/263) of strains, respectively, suggesting that *S. thermophilus* phages may carry anti-RM proteins.

**Table 1:**
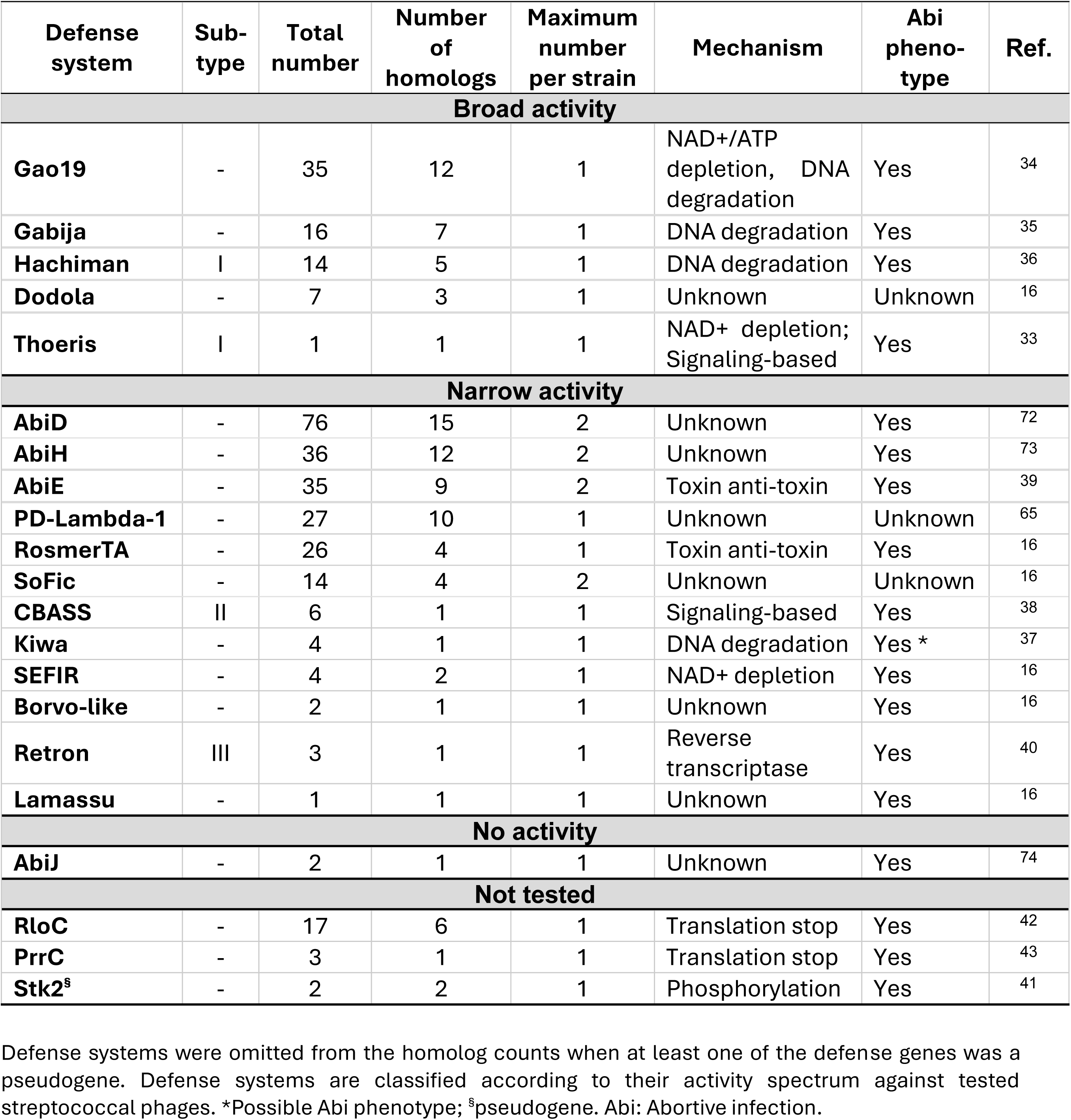
Information on additional defense systems found in *S. thermophilus*.

### Accessory defense systems provide resistance against dairy-associated phages

Eighteen defense systems were cloned and experimentally tested against 16 dairy phages, including 14 representatives from the five genera that infect *S. thermophilus,* as well as two phages (*Skunavirus* p2 and *Ceduovirus* c2) that infect *Lactococcus cremoris* (Fig. 3A). The efficacy of each defense system was evaluated by measuring the efficiency of plaquing (EOP) of phages on strains that harbor the plasmid-encoded systems. The efficacy of low-copy (pTRKL2) and high-copy (pNZ123 and pTRKH2) vectors^44,45^ were initially compared to identify the most suitable for assessing defense system effectiveness (Fig. 3B). Overall, pTRKL2 and pTRKH2 conferred higher levels of phage resistance, particularly with the Kiwa and Dodola defense systems. Notably, no phage resistance was observed for Dodola when the system was cloned into pNZ123, underscoring the critical role of plasmid selection in such experiments. This suggests that features like copy number and origin of replication significantly influence the outcomes. Based on these findings, we proceeded with the low-copy vector pTRKL2 for further cloning experiments.

**Fig. 3:**
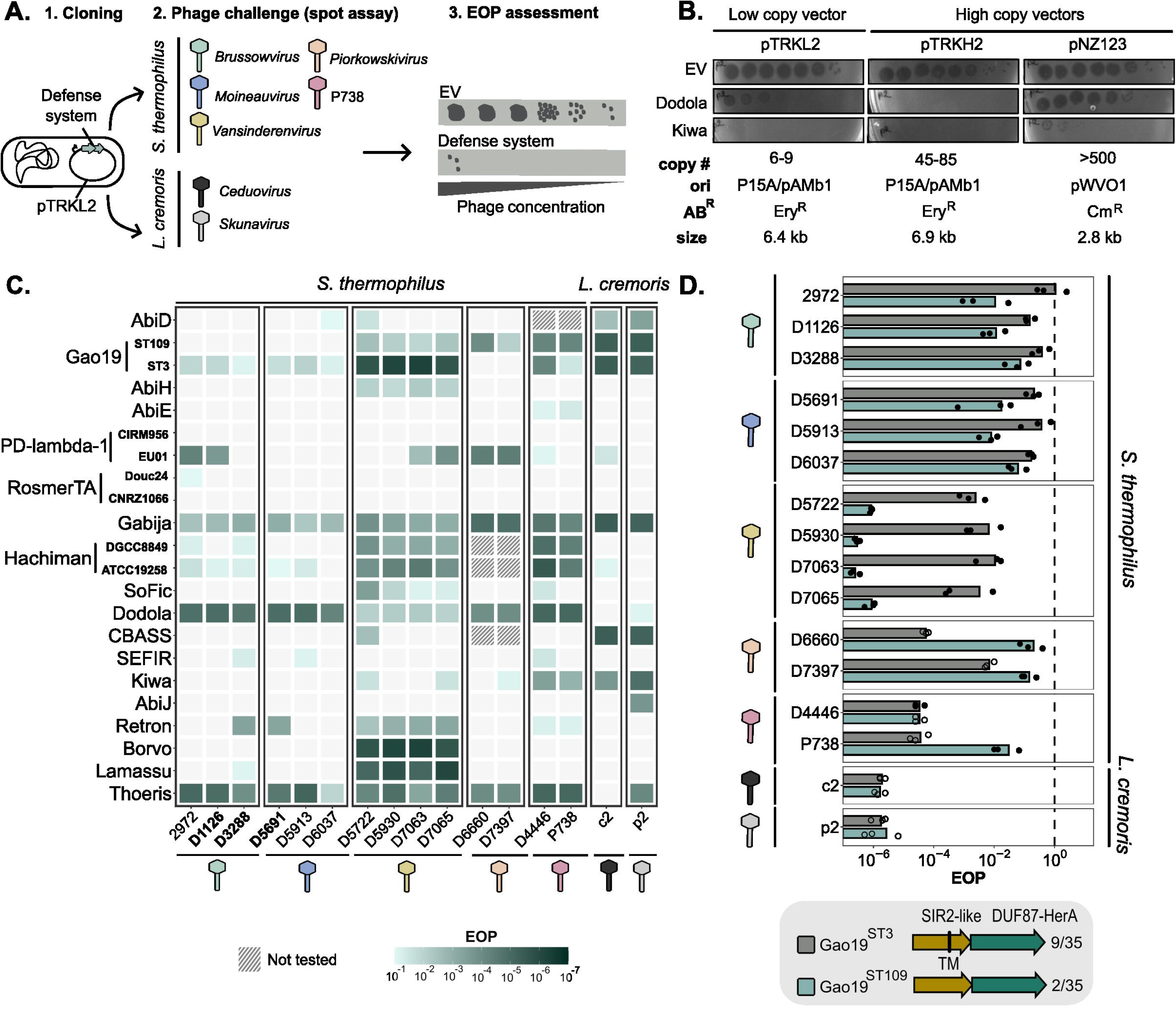
Activity of defense systems against phages infecting *S. thermophilus* and *L. cremoris.* **A.** Strategy for assessing the efficiency of several defense systems against streptococcal phages from five genera and lactococcal phages from two genera. Anti-phage activity was measured by evaluating the EOP of phages on the host strain containing the defense mechanism cloned on a plasmid vector compared to the host strain with the empty cloning vector. **B.** Comparison of Dodola and Kiwa defense systems efficiency against *L. cremoris* phage p2 when cloned into low-copy and high-copy vectors. **C.** Heatmap illustrating the EOP of phages on strains carrying each defense system. Darker shades indicate that the defense systems provided stronger anti-phage protection and an EOP of 10^-1^ was used as a minimum threshold for anti-phage activity. Three independent replicates were done. Some defense systems were not tested against few phages, as they are naturally present in the host strain. For the defense systems Gao19, PD-Lambda-1, RosmerTA, and Hachiman two homologs were tested. Phages encoding known ACR are highlighted in bold. The top to bottom presentation of the defense systems reflects the high to low prevalence. **D.** Comparison of anti-phage activity of two Gao19 homologs found in *S. thermophilus*. Gao19 was found in 35 strains, of which 9 corresponded to Gao19^ST3^ and 2 to Gao19^ST109^. Bars represent the mean EOP from three independent replicates. Filled circles indicate the presence of countable plaques, while hollow circles signify zones of lysis where plaques were not counted. AB^R^: Antibiotic resistance gene, Cm^R^: chloramphenicol resistance, DUF: Domain of unknown function; EOP: Efficiency of plaquing, Ery^R^: erythromycin resistance, EV: Empty vector, ori: Origin of replication, SIR: Sirtuin, TM: Transmembrane domain.

All tested defense systems demonstrated significant effectiveness, achieving at least 1-log reduction in titers against one or more of the tested phages (Fig. 3C). Specifically, 17 of the 18 tested defense systems were effective against streptococcal phages, with some providing protection up to almost 7 logs, thereby completely abolishing phage replication in certain cases (Supplementary Fig. S3). While defense systems such as Gabija, Dodola, Hachiman, and Thoeris conferred broad-spectrum protection across all tested streptococcal phages, other systems displayed more selective activity. For instance, AbiH and Borvo-like systems specifically inhibited the replication of *Vansinderenvirus* (Fig. 3C). Some defense systems, including SEFIR and RosmerTA, exhibited even narrower specificity, targeting only specific phages within a single viral genus. This underscores the importance of testing multiple phages, including closely-related ones, to uncover intra-genus specificities.

Of interest, *Brussowvirus* and *Moineauvirus*, which are the most prevalent phages in the industry, were less sensitive to the tested defense systems, with only four and three defense systems conferring some level of protection against all the phages from these viral genera, respectively. Overall, the level of protection was also higher against other genera (i.e., *Vansinderenvirus*, *Piorkowskivirus,* and P738), with most systems providing over a 3-log reduction in phage titers for these less common genera. *L. cremoris* phages c2 and p2 were also effectively targeted by these defense systems originating from *S. thermophilus,* with some offering high levels of protection.

Overall, the resistance patterns observed across our tested defense systems generally exhibited higher levels of protection compared to those reported by Kelleher *et al*^22^. While this difference may partly be explained by the use of different phages or defense homologs, it could also be attributed to our use of a low-copy vector plasmid, which likely enhanced the resistance conferred by systems like Gao19 and Gabija, where the same homologs were tested in both studies.

### Defense system homologs exhibit diverse activity spectra

Our bioinformatic analysis revealed that most defense systems had multiple homologs in *S. thermophilus* strains (Table 1). To explore whether these homologs offer different protection against our panel of phages, we conducted a comparative assessment of the activity of some of these homologs (Fig. 3C-D). The Gao19 defense system (also known as the SIR2-Her complex or Nezha) was identified in 35 strains, encompassing 12 homologs (Table 1). We compared the effectiveness of two of them, Gao19^ST3^ (present in nine strains) and Gao19^ST109^ (present in two strains). Despite sharing only 17.9% and 22% amino acid (aa) identity for the SIR2 and Her proteins, respectively, both homologs contained the same predicted domains (i.e., a SIR2-like protein and a DUF87 domain corresponding to a helicase domain HerA). However, the Gao19^ST3^ SIR2-like protein included a unique C-terminal transmembrane domain (Fig. 3D). We observed that while both homologs showed similar activity against most phages from the *Moineauvirus* and *Brussowvirus* genera, as well as those infecting *L. cremoris*, differences emerged with the other *S. thermophilus* phage genera. Gao19^ST109^ provided over 6-log protection against *Vansinderenvirus*, whereas Gao19^ST3^ achieved only a 2−3-log reduction, depending on the phage. Conversely, Gao19^ST3^ was more effective against *Piorkowskivirus* phages, providing protection with a reduction of 2−4 logs compared to 1 log for Gao19^ST109^.

Distantly related homologs of Hachiman (HamA and HamB subunits, sharing 15% and 17% aa identity, respectively) with similar domain annotations provided similar levels of resistance against the tested phages (Fig. 3C). In contrast, RosmerTA homologs, which were more distantly related (with toxins sharing 11% aa identity and antitoxins 6.7%), proved inefficient against most tested phages. The RosmerTA^CNRZ1066^ homolog lacked the peptidase M78 domain, which is typically found in this defense system. Interestingly, two homologs of the PD-Lambda-1 system, despite sharing 98.5% amino acid identity, displayed divergent anti-phage activity. PD-Lambda-1^CIRM956^ was completely ineffective, while PD-Lambda-1^EU01^ provided a high level of resistance (>3-log reduction) against some *Brussowvirus*, *Vansinderenvirus*, and *Piorkowskivirus* phages (Fig. 3C). This emphasizes the importance of testing different homologs to evaluate the specificity of defense systems, as even minor aa changes can significantly impact their activity.

### Defense systems can be integrated into the chromosome of an industrial strain

To effectively use a defense system against phages in industrial strains, it must be naturally transferable across strains via processes such as conjugation, transduction, or natural transformation. *S. thermophilus* can perform natural transformation, enabling it to uptake DNA, which can be leveraged to integrate a defense system into the chromosome^46^. This is achieved by transforming the strain of interest with a recombination template that contains a defense system and its native promoter, flanked by upstream and downstream homologous regions of the target locus (Fig. 4A). Transformed cells that have successfully integrated the defense system can then be screened by PCR following phage selection. Two streptococcal defense systems, PD-Lambda-1^EU01^ and Dodola, were selected for chromosomal integration into the industrial strain *S. thermophilus* DGCC7710.

**Fig. 4:**
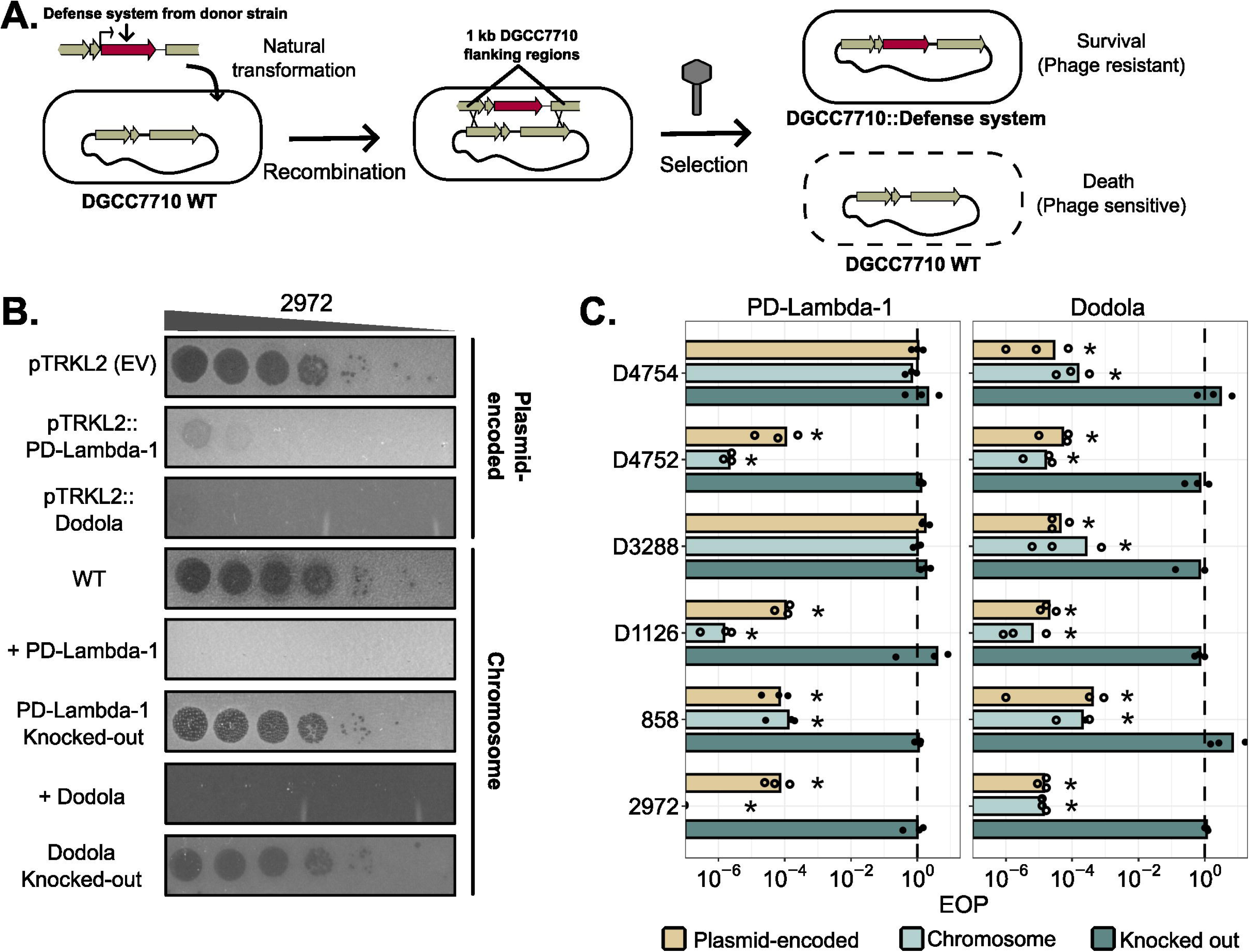
Enhancing *S. thermophilus* resistance through natural competence-mediated transformation of defense systems. **A.** Schematic illustration of the insertion of a defense system into the chromosome of *S. thermophilus* through natural competence. A recombination template, consisting of the defense system flanked by 1-kb regions that are homologous to the upstream and downstream sequences in the recipient strain (*e.g.* DGCC7710), was transformed. Phages were used to select bacteria that integrated the defense system. **B.** Spot test showing the comparison of defense-system (PD-Lambda-1 or Dodola) efficiency against phage 2972 when encoded on pTRKL2 vector versus when chromosomally integrated into DGCC7710. As a confirmation, the integrated defense systems were also knocked out of the chromosome. **C.** EOP of phages belonging to the *Brussowviruses* genus on strains carrying the PD-Lambda-1 and Dodola defense systems cloned on the pTRKL2 vector or integrated into the DGCC7710 chromosome. Each circle represents an individual replicate (N=3) with filled circles indicating the presence of countable plaques and hollow circles signifying zones of lysis where plaques were not observed. Asterisks indicate statistical significance (EOP different from 1) according to a t-test with a p-value less than 0.05. EOP: Efficiency of plaquing, EV: Empty vector, WT: Wild type.

PD-Lambda-1^EU01^ was integrated into the DGCC7710 genomic locus corresponding to that of the donor strain. Integrated PD-Lambda-1^EU01^ demonstrated robust effectiveness, providing protection against *Brussowvirus* phages comparable to that of the vector-encoded version (Fig. 4B-C). The lysis zone intensity at high phage concentrations was even reduced compared to the vector-encoded system (Fig. 4B). The role of PD-Lambda-1^EU01^ in phage resistance was further validated by the loss of protection when the integrated system was removed (Fig. 4B-C). The Dodola defense system was integrated into the same genomic locus as PD-Lambda-1, which is not where it naturally exists in *S. thermophilus* strains that encode this defense system. Nonetheless, DGCC7710 with the integrated Dodola showed high resistance against all *Brussowvirus* phages tested (Fig. 4B-C).

### Combining defense systems enhances immunity against phages

As shown in Fig. 1B, various defense systems often co-occurred within *S. thermophilus* strains, with an average of 7.5 systems per strain. Given the crucial role of the chromosomally-encoded CRISPR-Cas systems in generating resistant strains, we aimed to evaluate whether combining CRISPR-Cas with other defense systems could enhance protection against phages, particularly those encoding ACR proteins. A plasmid containing one defense system (Gabija, Hachiman, or Thoeris) was transformed into a CR1-immune DGCC7710 strain, which has a one spacer in its CR1 locus that targets the tested phages. The effectiveness of individual and combined defenses was assessed using solid and liquid assays (Fig. 5A and B).

**Fig. 5:**
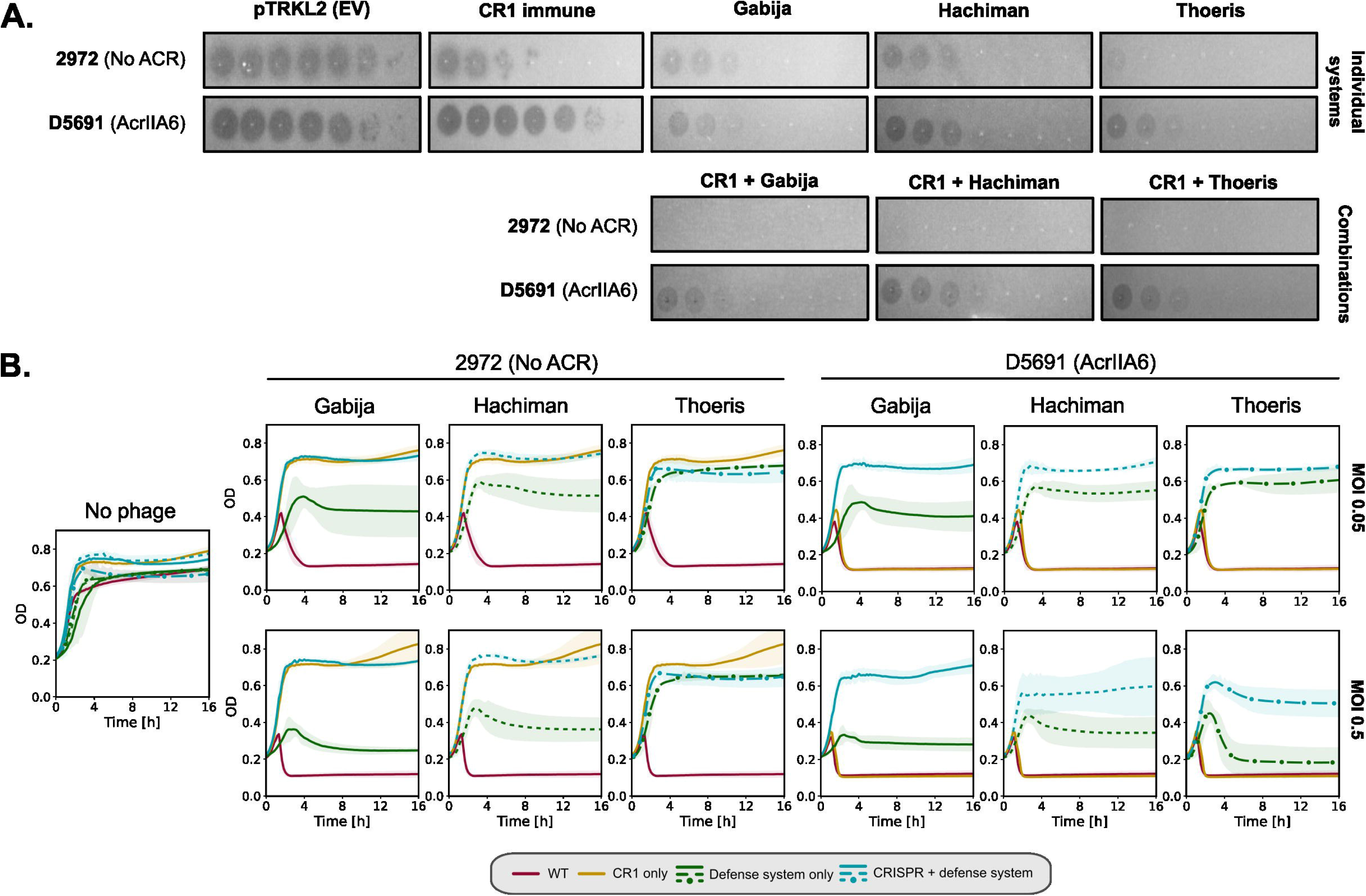
Combination of defense systems with CRISPR-Cas immunity improve phage resistance levels. Gabija, Hachiman, or Thoeris were transformed into a CR1-immune DGCC7710 strain, and their activity was compared to strains containing the individual defense systems alone. **A.** Examples of spot test results for phage D5691 encoding an AcrIIA6 with anti-CR1 activity and phage 2972 which does not carry known ACRs. **B.** Result of the liquid assay for the different defense system combinations with CR1 immunity at an MOI of 0.05. Control conditions (i.e., WT and CR1 only) correspond to DGCC7710 WT and CR1-immune DGCC7710 containing the empty pTRKL2 vector. Each line represents the mean of three independent replicates. The shaded areas represent confidence intervals. WT: Wild type.

CR1 immunity effectively protected against phage 2972, a phage that does not encode a known ACR protein, reducing its titer by 3 logs on M17 plates (Fig. 5A and Supplementary Fig. S4A). Non-CRISPR defense systems provided a 3−5-log protection against this phage when used individually (Fig. 5A and Supplementary Fig. S4A). Combining CR1 immunity with either Gabija, Hachiman, or Thoeris completely abolished phage replication, with no escape phages observed in spot tests (Fig. 5A). In liquid assays, combining CR1 with another defense system did not enhance resistance against phage 2972 at two multiplicities of infection (MOIs) (0.5 and 0.05) compared to CR1 alone, as the optical density (OD) curves showed the same trend and reached the same plateau (Fig. 5B). However, the assessment of the number of phages that were able to bypass both defense systems confirmed that compared to CR1 alone, no escape phage was present when an additional defense system was included (Supplementary Fig. S4B).

Phage D5691 encodes an AcrIIA6, which enables it to completely bypass the defense provided by the CR1 system (Fig. 5A-B and Supplementary Fig. S4A). However, D5691 was sensitive to Gabija, Hachiman, and Thoeris, each of which conferred a 3−4-log protection (Fig. 5A and Supplementary Fig. S4A). The improved protection when CR1 was combined with another defense system was particularly evident in liquid assays (Fig. 5B). For example, in the presence of phages at both MOIs, the expression of Gabija in CR1-immune DGCC7710 allowed bacterial growth to reach OD levels comparable to those observed in the absence of phages (Fig. 5B). The CR1-Gabija combination even demonstrated a synergistic effect, where the protection conferred by both systems together exceeded the sum of their individual contributions, as quantified by a synergy score (Supplementary Table S4). Similar trends were observed with Hachiman and Thoeris, though the synergy score was only statistically significant for an additive effect. In addition, the number of phage escape mutants was below the level of detection for all three 2-system combinations (Supplementary Fig. S4B).

Combinations of defense systems with CR3 immunity were also evaluated, demonstrating various levels of efficacy for different MOIs and phages (Supplementary Fig. S4B and S4C). Like the synergy observed with the CR1-Gabija combination, CR3 and Gabija also demonstrated synergistic effects (Supplementary Table S4). The ability to target phages carrying AcrIIA3 (encoded by D5691) and AcrIIA5 (encoded by D1126) was also shown (Supplementary Fig. S4A). Additionally, using phage D3288 (which was not targeted by the spacers of the CR1 and CR3 arrays), we showed that combining an additional defense system can neutralize phages that would otherwise evade CRISPR defenses due to the absence of a matching protospacer (Supplementary Fig. S4A).

## 3. Discussion

Preventing phage infection remains an industrial challenge. A primary strategy to control these infections involves optimizing bacterial cultures by exposing strains to phages and naturally selecting BIMs with enhanced resistance. In *S. thermophilus*, CRISPR-Cas systems are the primary drivers of BIM selection through the acquisition of novel spacers in their CR1 or CR3 arrays, as very few non-CRISPR BIMs have been described for this bacterium^47,48^. While CRISPR-Cas remains the most effective tool for developing phage-resistant *S. thermophilus* strains, phages can overcome this resistance by mutating their protospacers or PAM^10^. Approximately 40% of streptococcal phages were also found to encode at least one known ACR protein, enabling them to bypass CRISPR type II-A systems. This limitation necessitates a search for alternative defense mechanisms to develop more effective strategies for combating streptococcal phages in the dairy industry.

In many other bacteria, the primary defense mechanism against phages is the alteration of the receptor required for phage adsorption. While *S. thermophilus* BIMs with mutations in genes involved in the biosynthesis of the polysaccharide receptor have been isolated, they exhibit an altered growth phenotype that hinders their industrial use^49^. Bacteria can carry a multitude of other intracellular defense systems capable of interrupting the viral cycle at every stage, from genome injection to host lysis^4^. Recent advances in predictive tools for identifying known defense systems in microbial genomes uncovered that on average, bacteria encode approximately five such systems^23^. This suggested that *S. thermophilus* likely harbors additional defense mechanisms beyond the CRISPR-Cas systems. Our analysis revealed that strains encode an average of 7.5 defense systems, with CRISPR-Cas and RM systems being the most prevalent. This higher prevalence of defense systems likely underlines their necessity for *S. thermophilus* to thrive in a changing virus-containing ecosystem.

Here, we identified 21 accessory defense systems, each present in less than 30% of the analyzed strains, significantly expanding the defense repertoire of *S. thermophilus*. Interestingly, most of these accessory systems have been shown to cause the collapse of the bacterial population at high MOI, a phenotype commonly associated with Abi systems that prevents the viral outbreak from spreading to neighboring cells^50^. We speculate that *S. thermophilus* may have evolved various Abi systems as a secondary line of defense, which is activated when phages have managed to circumvent CRISPR-Cas and/or RM systems. Abi systems such as PrrC and RloC, detected in several *S. thermophilus* strains, are known to be triggered when phage proteins inactivate type I RM systems^42,43^.

Seventeen defense systems demonstrated variable levels of resistance against a diverse panel of streptococcal phages. Among these, *Moineauvirus* and *Brussowvirus,* the two most frequently encountered genera in the industry, were the least susceptible to the tested defense systems. In contrast, other less commonly isolated viral genera displayed greater sensitivity. These rarer genera, which share homologies with lactococcal and non-dairy streptococcal phages, are thought to have emerged more recently than *Moineauvirus* and *Brussowvirus*, which have been isolated for decades^51–53^. This suggests that the latter may have evolved multiple strategies to evade *S. thermophilus* defense systems, contributing to their success in the dairy environment. Notably, some defense systems are activated when they sense the presence of particular phage proteins. When phages have mutations in these activators, or if they are absent, they can bypass the bacterial defense mechanisms^54^. Structural proteins, in particular, have been implicated in the activation of some defense systems^55^. Comparative genomics of the structural modules in *S. thermophilus* phages revealed significant divergence, potentially explaining the differences in sensitivity among certain phages^56^. Additionally, phages can escape defense systems by encoding counter-defense proteins^57^. Our analysis showed that ACR proteins are more prevalent in *Moineauvirus* and *Brussowvirus*, and other, not yet identified, anti-defense proteins may also be present. However, the absence of significant candidates predicted by bioinformatics tools suggests that these anti-defense proteins may differ from those known so far, which have predominantly been identified in phages infecting Gram-negative bacteria^58^.

Transferring these anti-phage systems into industrial strains is crucial for generating phage-resistant derivatives for food applications, but this process requires careful consideration as the use of genetically engineered organisms has generally a negative perception to consumers^59,60^. The European regulatory authorities do not classify strains as genetically modified organisms (GMOs) when genes are mobilized via transduction, conjugation, or transformation, provided the genetic material originates from the same species as that of the recipient strain^59^. As most anti-phage systems identified in *S. thermophilus* are chromosomally encoded, the preferred method for integrating the desired defense system is to exploit natural transformation. We successfully mobilized PD-Lambda-1 and Dodola into a specific genomic locus of an industrial strain. The resulting chromosomally-encoded systems provided robust phage resistance, surpassing even the abilities of plasmid-encoded versions. These results demonstrate the potential of this approach for developing phage-resistant strains within the constraints of dairy industry regulations.

Given the reliance on CRISPR-Cas systems for phage-resistance purposes in *S. thermophilus*, new defense mechanisms should be used in combination with CRISPR-Cas rather than replacing them. Previous research has demonstrated that CRISPR-Cas systems are compatible with type II RM systems, offering additive protection against streptococcal phages^61^. Our study further demonstrates that other DNA-degrading systems, such as Gabija and Hachiman, as well as NAD+ depletion-based systems like Thoeris, are also compatible with CRISPR-Cas, leading to increased resistance against phages, particularly those carrying ACR proteins. It is anticipated that additional defense systems will exhibit similar compatibility with CRISPR-Cas, as defense mechanisms generally work well together and can even exhibit synergistic effects^62^. This synergy was clearly observed when we combined Gabija and CRISPR-Cas systems. Moreover, these additional defense systems enabled the targeting of CEMs, significantly reducing the number of phages. This pyramiding strategy, where multiple defense systems are stacked within the same strain, has proven more effective at preventing the emergence of escape mutants compared to the mixing strategy, where strains with individual defense systems are combined^63^. While it is expected that stacking more defense systems will result in stronger phage resistance, it is important to consider the potential fitness costs associated with maintaining these systems^64^. It is crucial to ensure that key industrial traits for fermentation, such as acidification rates and growth rates, remain unaffected in these enhanced strains.

In conclusion, developing phage-resistant *S. thermophilus* strains is crucial for the success and sustainability of the milk fermentation industry. Our research highlights the importance of understanding and leveraging the diverse defense mechanisms inherent to this bacterium. By strategically combining CRISPR-Cas systems with other defense systems, we can create a robust and multi-layered defense to combat phages. This comprehensive strategy promises to enhance the resilience of starter strains, ensuring the high quality of fermented dairy products.

## 4. Materials and Methods

### Defense system prediction

*S. thermophilus* genomes (N=263) were downloaded from the NCBI Reference Sequence (RefSeq) database in March 2023 (Supplementary Table S1). Defense systems were predicted using PADLOC^24^ v2.0.0 and DefenseFinder^23^ v1.2.2. Predictions labelled as “other”, “cas_adaptation”, “cas_cluster”, or candidate systems were excluded as they did not represent complete or experimentally validated systems. Results from both prediction tools were combined using an in-house script. After manual inspection, the defense systems Viperin_solo and PD-T4-6 were excluded from the analysis as they appeared to be wrong predictions. The same genes that matched the closely related PD-T7-2 and Gao19 systems^65^ were combined into a single system denoted as Gao19. Similarly, Abi2 and AbiD, which mostly match the same genes, were clustered as AbiD systems. Type II and type IIG RM systems were grouped into RM type II. One CRISPR II-A was incorrectly predicted as CRISPR II-C and was manually corrected. The detailed list of identified defense systems can be found in Supplementary Tables 2 and 3. A core proteome species-phylogeny was generated with OrthoFinder^66^.

### Plasmid and prophage predictions

Plasmids were identified in incomplete genomes using the PlasmidFinder database with Abricate v1.0.0 (https://github.com/tseemann/abricate). Prophages were identified with Phigaro v2.4.0, and hits greater than 10 kb were confirmed using PHASTEST webserver^67^.

### Restriction site analysis

RM proteins were submitted to the REBASE database (http://rebase.neb.com/rebase/) to predict restriction sites (identity threshold of 75%). For type I, only the specificity protein was used whereas both the methylase and endonucleases were used for types II and III. Detailed information for each RM system is provided in Supplementary Table S5. To evaluate the avoidance of restriction sites, the genomes of 191 phages (116 *Moineauvirus*, 46 *Brussowvirus*, 14 *Piorkowskivirus*, 13 *Vansinderenvirus,* and 2 P738-like phages) that infect *S. thermophilus* were downloaded from NCBI in June 2023 (Supplementary Table S6). Predicted sites for types II and III RMs were counted in each genome (i.e., observed number of restriction sites). The expected number of restriction sites was estimated using a Markov-immediate neighbor dependence model^68^. Briefly, the frequency of each nucleotide and dinucleotide was calculated for each genome. For example, the frequency for GATC (𝑓_𝐺𝐴𝑇𝐶_), was calculated as:

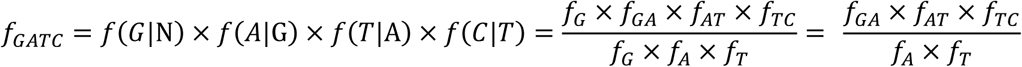

where N represents any nucleotide, and, for example, 𝑓(𝐴|G) is the frequency of observing an A preceded by a G.

The expected number of restriction sites in each genome was obtained by multiplying the corresponding frequency by the genome length. Restriction site avoidance was calculated using the ratio between the observed and expected numbers of sites. Underrepresented sites had ratios < 0.75, and overrepresented sites had ratios > 1.25.

### ACR and methyltransferase prediction in phage genomes

ACRs proteins were predicted by dbAPIS^58^ with a threshold for e-value relevance set at e^-30^. Methyltransferases were identified by retrieving phage proteins with annotations containing “methyltransferase” or “methylase” and confirming their methyltransferase function using REBASE database and InterPro^69^.

### Bacterial growth conditions

Bacteria and phages used in this study are listed in Supplementary Table S7. All strains were grown overnight unless stated otherwise. *Escherichia coli* was grown at 37°C with shaking in Brain Heart Infusion (BHI, Difco) medium. *S. thermophilus* was grown in 0.5x M17 (Nutri-Bact) supplemented with 0.5% w/v lactose (LM17) and incubated at 37°C without shaking. *L. cremoris* was grown in M17 with 0.5% w/v glucose (GM17) and incubated at 30°C without shaking. For solid media, 1.5% agar (Laboratoire Mat) was added, and plates were incubated at 37°C for *E. coli*, 42°C for *S. thermophilus*, and 30°C for *L. cremoris*. Erythromycin was added to the media, when necessary, to maintain pTRKL2 or pTRKH2 vectors, at final concentrations of 150 µg/ml for *E. coli* and 5 µg/ml for *S. thermophilus* and *L. cremoris*. Chloramphenicol was used at final concentrations of 20 µg/ml and 5 µg/ml to maintain pNZ123 vector in *E. coli* and *L. cremoris*, respectively.

### Construction of defense system-encoding plasmids

Some of the examined defense systems were PCR amplified from our bacterial collection using primers listed in Supplementary Table S8 using the Q5 High-Fidelity DNA Polymerase (New England Biolabs). Other defense systems were synthesized at Bio Basic or Integrated DNA Technologies. Each defense system was then cloned into the low copy vector pTRKL2^44^ (and high-copy vector pNZ123^45^ for Kiwa and Dodola) via Gibson assembly and transformed into chemically competent *E. coli* NEB5α. PCR screenings were performed with Taq DNA polymerase (Bio Basic), positive clones were verified by Sanger sequencing, and plasmids were extracted using the QIAprep Spin Miniprep Kit (Qiagen). Plasmids were then electroporated into *L. cremoris* MG1363^70^. For *S. thermophilus*, most plasmids were electroporated into relevant strains using the protocol described previously^71^ with the following modifications: after the 2-h incubation at 42°C, the transformation mix was transferred to 10 ml LM17 supplemented with erythromycin, incubated overnight at 42°C, and plated the following day. For plasmids/strains that do not readily electroporate, plasmids were introduced by natural competence using a protocol described elsewhere^46^, except that competent cells were grown in milk. Prior to transformation, we made sure that the tested defense system was not naturally present in the bacterial strains using Padloc and DefenseFinder. The correct sequences for the defense systems found in the transformants were verified by Sanger sequencing.

To evaluate the combinations of other defense systems with CRISPR-Cas, plasmids encoding Gabija, Hachiman, or Thoeris were transformed into either CR1-immune DGCC7710, whose CR1 locus contained a spacer (5’-AAGTAGCCATACAAGAAGATGGATCAGCA-3’) targeting *orf20* of phage 2972 and its homologs, or into CR3-immune DGCC7710, whose CR3 locus contained a spacer (5’-CTGATGGAACCTGGCCACTGCAACCACGAC-3’) targeting the same *orf20* and its homologs.

### Assessment of defense system efficiency

The effectiveness of each defense system was evaluated through spot tests using phage lysates with titers of *ca.* 10^8^–10^9^ PFU/ml. Briefly, molten agar was mixed with 15% of overnight culture and poured onto 1% agar M17 plates containing 10 mM CaCl_2_. Next, 3 μl drops of serially diluted (tenfold) lysate were spotted onto the plates and let dry before incubation. For *S. thermophilus*, the molten agar contained LM17, 0.5% glycine, 0.4% agarose, and 0.1% milk. For *L. cremoris*, the molten agar was made of GM17, 0.75% agar, 10 mM CaCl_2_. Phage titers were evaluated after overnight incubation. The efficiency of plaquing (EOP) was calculated as the ratio of the phage titer in the presence of the defense system to that obtained with the strain carrying the empty vector.

### Chromosomal insertion of defense systems

PD-Lambda-1 or Dodola system was inserted between genes CW339_RS00280 and CW339_RS00285 in the chromosome of *S. thermophilus* DGCC7710. PD-Lambda-1, including its native promoter, was amplified from the donor strain *S. thermophilus* DGCC688. Additionally, 1-kb upstream and downstream flanking regions of the insertion site in DGCC7710 were amplified by PCR. The recombination template was constructed by Gibson assembly of the three fragments followed by PCR amplification with external primers (Supplementary Table S8). For Dodola (*S. thermophilus* TK-P3A), the recombination template was synthesized at GeneScript. Then, 10 μg of recombination template were transformed via natural competence into DGCC7710. The transformation mix was added to LM17 and incubated at 42°C. The following day, 500 μl of an overnight culture were mixed with 3 ml molten agar and 100 μl of phages D1126 and 2972 (*ca.* 10^8^ PFU/ml) to select for cells that had integrated PD-Lambda-1. For Dodola, phage D5691 (*ca.* 10^8^ PFU/ml) was used to select transformants. The genome insertion and the absence of new spacers in the CR1 and CR3 loci were verified by PCR and Sanger sequencing. To confirm the observed anti-phage activity, the defense system was also knocked out from the transformants (DGCC7710::defense_system_chrs_) using a recombination template containing only the upstream and downstream flanking regions. Knockout mutants were confirmed via PCR screening.

### Killing curve assay

Overnight cultures were adjusted to an OD of 0.2 (*ca*. 10^7^ CFU/ml) in LM17 supplemented with 10 mM CaCl_2_ and 180 μl were transferred to each well of a 96-well plate. Twenty microliters of buffer or lysate (phage 2972 or D5691) were diluted to obtain a MOI of 0.5 or 0.05. Plates were incubated for 16h at 37°C, with OD_600_ measurements taken every 10 min after orbital shaking for 10 seconds. Each curve represents the mean of three biological replicates. After 24h, the plate was centrifuged (4000g, 5min), and 100 μl of the supernatant was recovered, diluted, and spotted to assess the titer of escape mutants. Synergy between combined defense systems was evaluated by calculating the difference between the OD of the combination and the sum of the OD values for the CRISPR defense alone and the additional defense system alone. A positive score indicates a synergistic effect, a null score reflects an additive effect of the two defenses and a negative score indicates that the combined systems do not provide enhanced resistance compared to either system used alone (mostly CRISPR). Synergy scores were calculated from three independent replicates, and a t-test was performed to assess statistical significance.

## 7. Acknowledgements

The authors extend their gratitude to Vincent Somerville for assistance with phylogenetic tree generation, to Carlee Morency for help with the AbiD experiments, and to Anne Millen for discussions. Additionally, we acknowledge the support of two FRQNT networks, PROTEO and Op+Lait, for providing the opportunity to present this work at international conferences. We also thank Crayon-Bleu for editorial assistance.

## 8. Author contributions

A.L., D.A.R., and S.M. conceived the study. A.L., G.R., and S.M. designed the experiments. A.L. and J.L. conducted the experiments. D.M. and P.H. provided feedback on experiments. A.L. performed data analysis and wrote the first draft of the manuscript. All authors edited and approved of the manuscript.

---

**Fig. S1:**
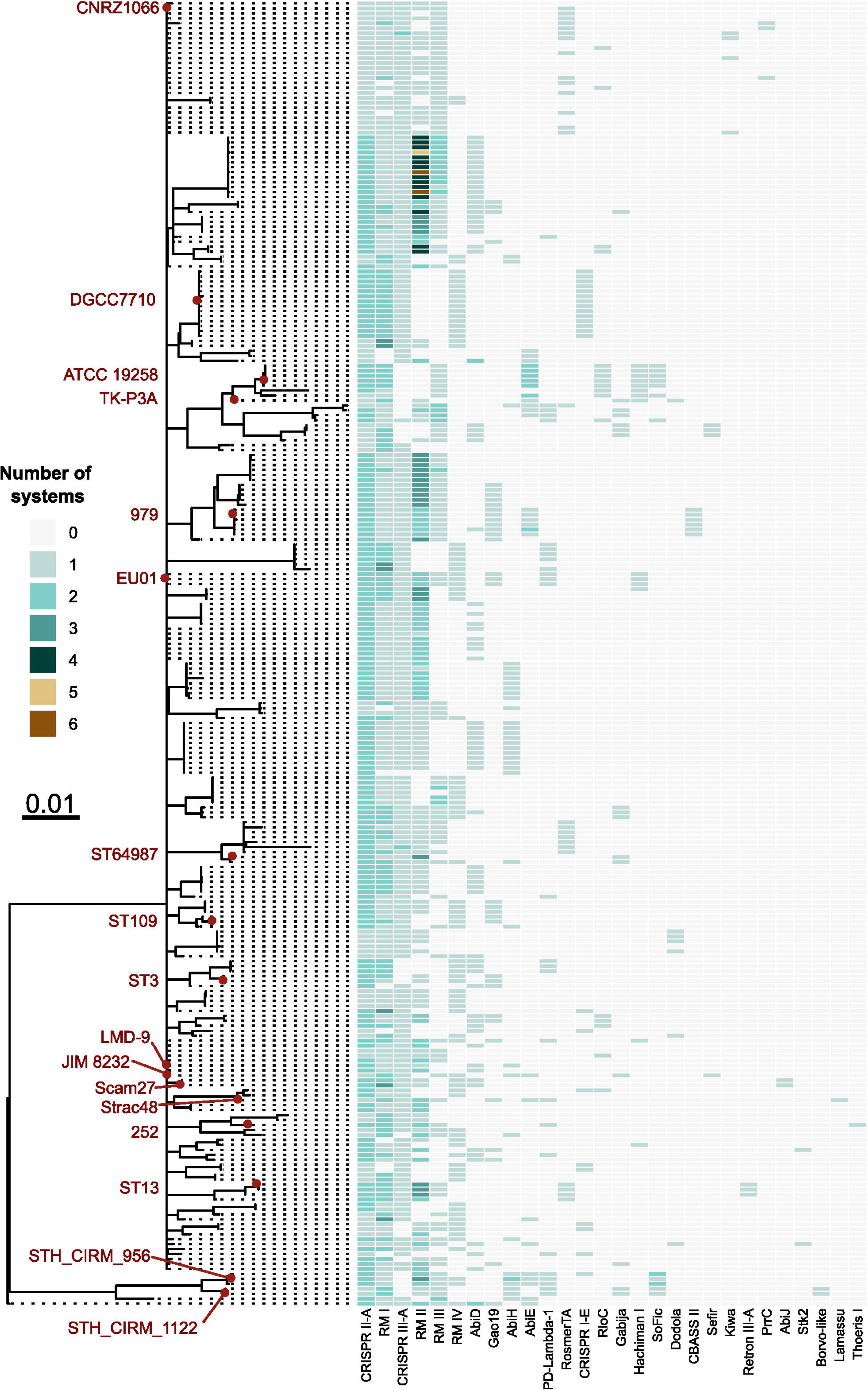
Phylogenetic tree of *S. thermophilus* genomes and their defense systems. The heatmap indicates the presence and number of defense systems within each genome. Strains with defense systems that were used in the experimental part of this study are highlighted in red. For additional details, refer to Tables S3 and S7.

**Fig. S2:**
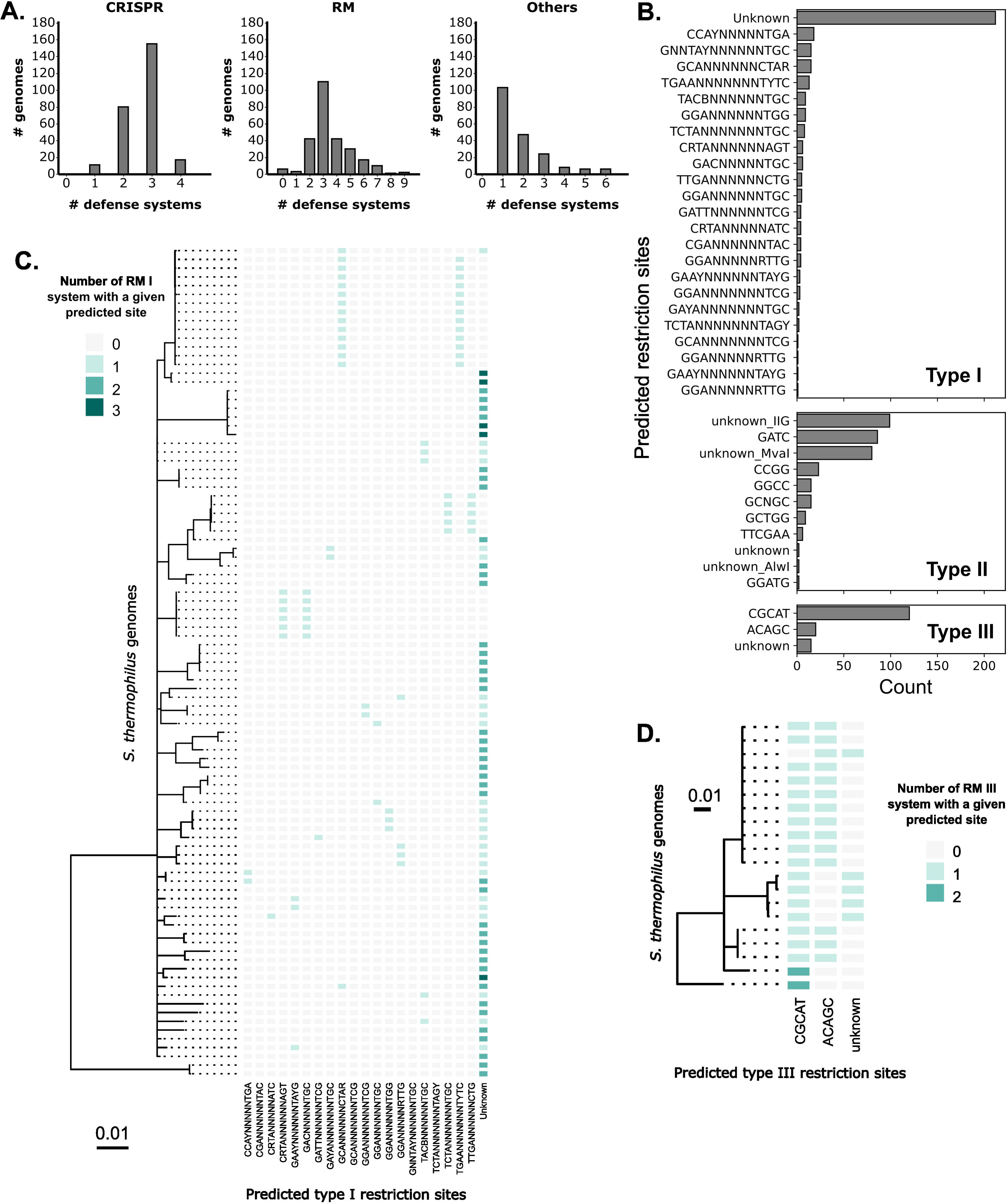
Additional information on the prevalence of defense systems and RM-predicted recognition sites. **A.** Histograms showing the number of defense systems per strain according to the categories CRISPR, RM, and other systems. **B.** Prevalence of each predicted restriction site according to the RM type. For type II systems, the category “unknown” was divided into “unknown_MvaII”, “unknown_AlwI”, “unknown_IIG”, and “unknown” based on the restriction enzyme annotation (MvaI/BcnI family or AlwI family) or RM type II (type IIG). **C-D.** Heatmaps displaying co-occurrence within the same genome of RM type I (**C.**) and type III systems (**D.**) with different predicted restriction sites. Only *S. thermophilus* strains with more than two RM type I or RM type II systems are shown and represented on the phylogenetic tree.

**Fig. S3:**
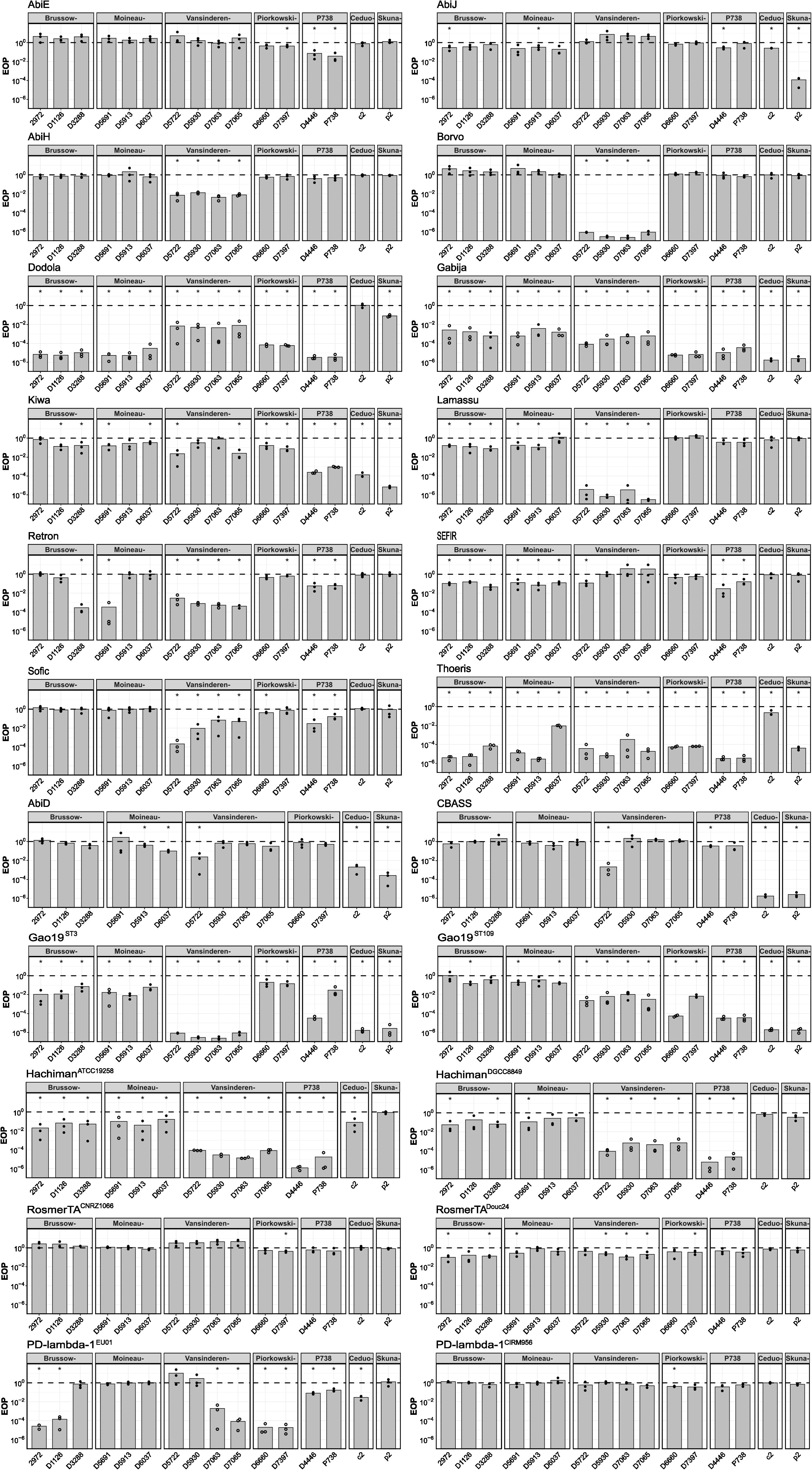
Detailed spot test results for each tested defense system using streptococcal and lactococcal phages. Bars represent the mean EOP from three independent replicates which are each represented by a circle. Filled circles indicate the presence of countable plaques, while hollow circles signify zones of lysis where plaques were not observed. The dotted line represents an EOP of 1, indicating no phage defense. Asterisks indicate statistical significance (EOP different from 1) according to a t-test with a p-value less than 0.05. Hyphen on top of each graph replaces the word *virus*.

**Fig. S4:**
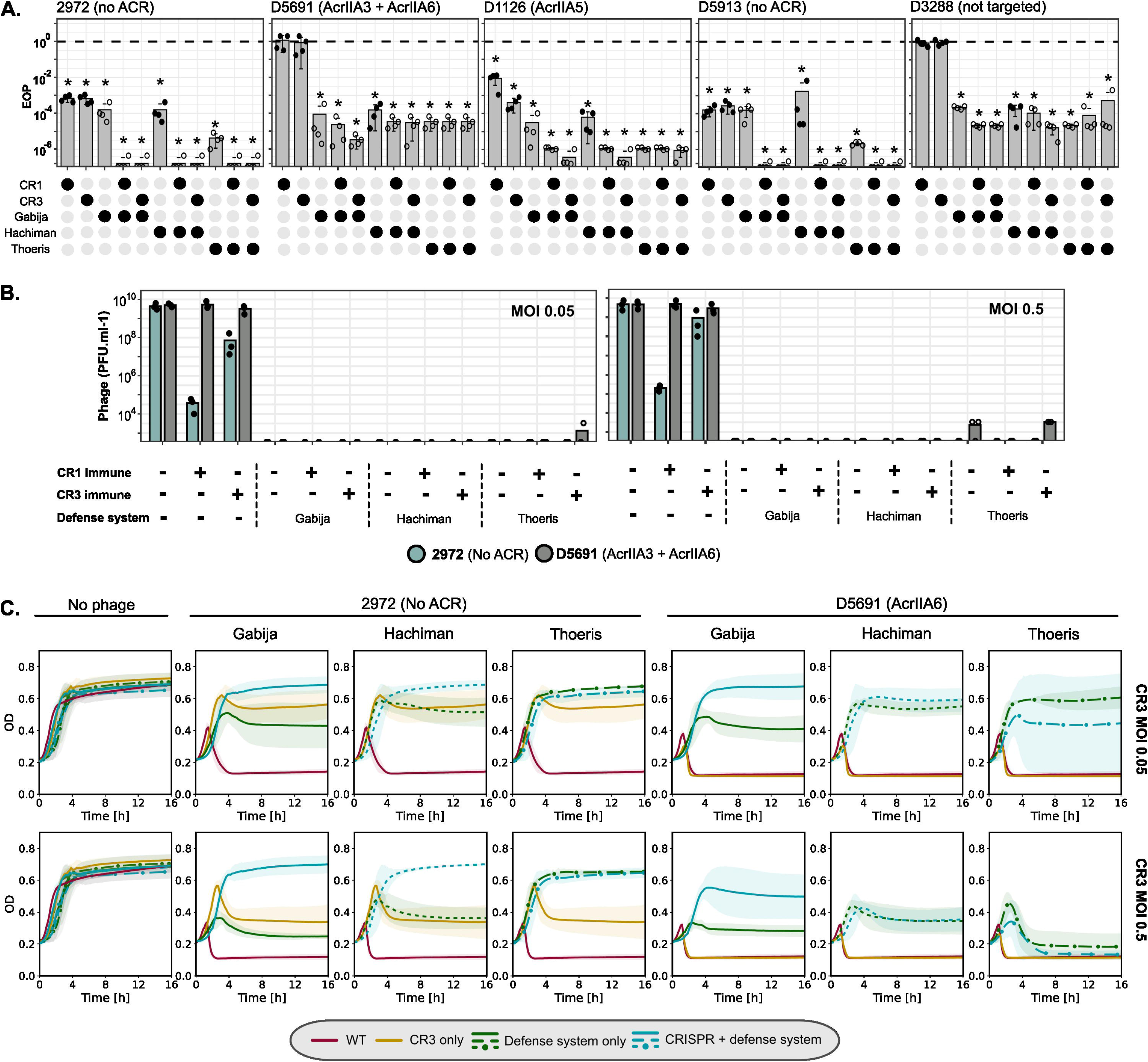
Detailed results of phage resistance using various combinations of CRISPR-Cas systems with additional defense systems. **A.** Detailed spot test results for combinations of Gabija, Hachiman, or Thoeris with either CR1 or CR3 immunity. Five phages were tested: D5691 which encodes an AcrIIA3 (anti-CR3 activity) and AcRIIA6 (anti-CR1 activity), D1126 which encodes an AcrIIA5 (anti-CR1 and anti-CR3 activities), 2972 and D5913 which do not carry known ACRs, and D3288 which is not targeted by the spacers in the CR1- and CR3-immune strains. Bars represent the mean EOP from three independent replicates, which are each represented by a circle. Filled circles indicate the presence of countable plaques, while hollow circles signify zones of lysis where plaques were not observed. The dotted line represents an EOP of 1, indicating no phage resistance. Asterisks indicate statistical significance (EOP different from 1) according to a t-test with a p-value less than 0.05. **B.** Evaluation of the number of plaque forming units/escaping phages after the liquid assays for phages 2972 and D5691 depending on the MOI. **C.** Liquid assays for the phage defense combinations with CR3-immune DGCC7710 at MOI 0.5 and 0.05. Each line represents the mean of three independent replicates. The shaded areas represent confidence intervals.

